# Structural Insights of SARS-CoV-2 Spike Protein from Delta and Omicron Variants

**DOI:** 10.1101/2021.12.08.471777

**Authors:** Ali Sadek, David Zaha, Mahmoud Salama Ahmed

**Author notes:** Corresponding author: **Mahmoud S. Ahmed, PhD**, Department of Internal Medicine/Division of Cardiology, University of Texas Southwestern Medical Center, 6000 Harry Hines Blvd, Dallas, Texas, 75390, USA. Authors contributed equally (Ali Sadek, David Zaha).

## Abstract

Given the continuing heavy toll of the COVID-19 pandemic and the emergence of the Delta (B.1.617.2) and Omicron (B.1.1.529) variants, the WHO declared both as variants of concern (VOC). There are valid concerns that the latest Omicron variant might have increased infectivity and pathogenicity. In addition, the sheer number of S protein mutations in the Omicron variant raise concerns of potential immune evasion and resistance to therapeutics such as monoclonal antibodies. However, structural insights that underpin the potential increased pathogenicity are unknown. Here we adopted an artificial intelligence (AI)-based approach to predict the structural changes induced by mutations of the Delta and Omicron variants in the spike (S) protein using Alphafold. This was followed by docking the human angiotensin-converting enzyme 2 (ACE2) with the predicted S proteins for Wuhan-Hu-1, Delta, and Omicron variants. Our in-silico structural analysis indicates that S protein for Omicron variant has a higher binding affinity to ACE-2 receptor, compared to Wuhan-Hu-1 and Delta variants. In addition, the recognition sites of the receptor binding domains for Delta and Omicron variants showed lower electronegativity compared to Wuhan-Hu-1. Importantly, further molecular insights revealed significant changes induced at fusion protein (FP) site, which may mediate enhanced viral entry. These results represent the first computational analysis of structural changes associated with Omicron variant using Alphafold, Collectively, our results highlight potential structural basis for enhanced pathogenicity of the Omicron variant, however further validation using X-ray crystallography and cryo-EM are warranted.

## Introduction

In late December 2019, SARS-CoV-2 was identified as the etiological agent of a severe acute respiratory disease named as COVID-19 at Wuhan, China [1-3]. Since then, the virus propagated worldwide, causing an unprecedented magnitude of morbidity and mortality. Despite efforts by the scientific community, which yielded a number of highly effective vaccines and antiviral agents, the virus was able to undergo a number of mutations leading to the infection of 266 million reported cases and death of 5.26 million patients worldwide [4]. SARS-CoV-2 is a betacoronavirus and its genome encodes several structural proteins, including the glycosylated spike (S) protein that mediates viral attachment to the cells by binding to angiotensin-converting enzyme 2 (ACE2), a membrane bound carboxypeptidase [5-7]. Viral entry also appears to be mediated by priming of S protein facilitated by the host cell-produced serine protease TMPRSS2 [5, 8]. In addition, the viral genome also encodes nonstructural proteins including an RNA-dependent RNA polymerase (RdRp), main protease (M^pro^), and papain-like protease (PLpro) [9-11].

Although the viral evolution is slowed by the RNA proofreading capability of its replication machinery [12], several variants of interest (VOI) or concern (VOC) were discovered at the S protein surface including Alpha, Beta, Gamma, Delta, Lambda, Mu, and most recently Omicron [13-17]. Delta (B.1.617.2) and Omicron (B.1.1.529) variants showed high infection and transmission rates starting from late 2020 till late 2021 worldwide, where the WHO changed their status from VOI to VOC. This raises the concerns about the efficiency of existing vaccines to mitigate the emerging mutations, which prompted a campaign to deliver booster doses to enhance immunity in the face of the emerging variants. Both the Delta and Omicron variants appear to have undergone several mutations in the S-protein, which highlights the importance of better understanding of the structural consequences of these mutations to inform the development of therapeutics and vaccines. Interestingly, the mutations at S protein were not limited to the receptor-binding domain (RBD) in S1 where the viral receptor angiotensin-converting enzyme 2 (ACE2) binds, but extended to include N-terminal domain (NTD), S1/S2’ cleavage site, and fusion peptide (FP) including deletions, insertions and modification of affected amino acids [18]. Several structural studies have elegantly demonstrated the conformational dynamics of trimeric S-glycoprotein w/o ACE2 receptor in the closed (pre fusion) and open (post fusion) conformers [14, 19-21]. In addition, the D614G mutation, where ASP614 (D614) was changed to GLY, a common mutation in most of the newly identified variants was detected and structurally resolved using cryo-ET[18, 21, 22]. The Omicron variant has >30 mutations at different sites of the S protein, however the full structure for the entire S protein carrying these mutations is yet to be revealed. Therefore, the purpose of our study is to predict the structure of the entire S protein of the Delta and Omicron variants using the cutting-edge AI-enabled protein structural prediction tool, Alphafold [23]. This is followed by protein-protein docking of ACE2 receptor with the S protein of Wuhan-Hu-1, Delta, and Omicron strains using Clustpro [24].

## Methods

### Data Access

The FASTA amino acids sequence of the S protein of SARS-CoV-2 strain without mutations was retrieved from Uniport (Access no: P0DTC2). The amino acid sequences of the spike protein of Delta and Omicron variants were obtained from GSAID [25].

### Structure prediction using Alphafold

The primary amino acid sequences were submitted to AlphaFold2 to analyze the effects of recent mutations on the S proteins of the Delta and Omicron Variants, versus Wuhan-Hu-1. All calculations were conducted using a high-performance Web GUI GPU node on the UTSW BioHPC with 16 cores. We used the full dBs preset with model 2 for full computational flow. After completion, all the predicted structures enrolled AMBER force field relaxation phase to avoid any stereochemical violations, where the relaxed .pdb files were further analyzed and the PKL files were processed to deduce the predicted local distance difference test (pLDDT) values for each residue.

### Protein-Protein Molecular Modeling and Scoring

The three predicted structures of the whole S protein (Wuhan-Hu-1, Delta, and Omicron) strains were docked to ACE-2 receptor (PDB ID: 6M0J, chain A [26]) using the Clustpro protein docking server (https://cluspro.org/login.php) [24]. The whole energy profiles were evaluated based on the binding interaction of ACE-2 and the RBD of S protein using electrostatic and Van der Waals interactions (vdW), applying the following equation.

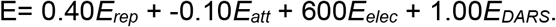

The top models were selected for generation of electrostatic potential map and visualization using Pymol (version 2.0.6 by Schrödinger).

## Results and Discussion

### Structural Characterization of S protein of Wuhan-Hu-1 strain and ACE-2

We validated the S protein structure using Alphafold to reveal the electrostatic map of the receptor binding domain for WT and the overall structure prior to application of mutations (mean pLDDT = 77.83) with an overall of 1273 amino acids (**figure 1**). The electrostatic map showed the abundance of electronegativity at the receptor binding domain of S protein where ACE-2 receptor binds (**figure 2**). This was followed by docking of S protein to the ACE-2 receptor extracted from PDP (PDB ID: 6M0J) using Clustpro to validate the previously resolved recognition sites for binding of ACE-2 receptor to the RBD of S protein, which highlighted the importance of T470-T478 loop and Y505 at ΔG = -250.20 Kcal/mol based on the electrostatic and vdW scoring functions [27]. One of the interesting amino acid residues is D614 from Sub-Domain 2 that shows hydrogen bonds along with K835, Y837, and K854 of fusion protein, which suggests the importance of D614 in the stabilization of FP [28, 29]. Based on our docking studies, we demonstrate the proximity of D614 along with FP of S-protein, confirming its potential role to stabilize the FP (**figure 3C**).

**Figure 1.**
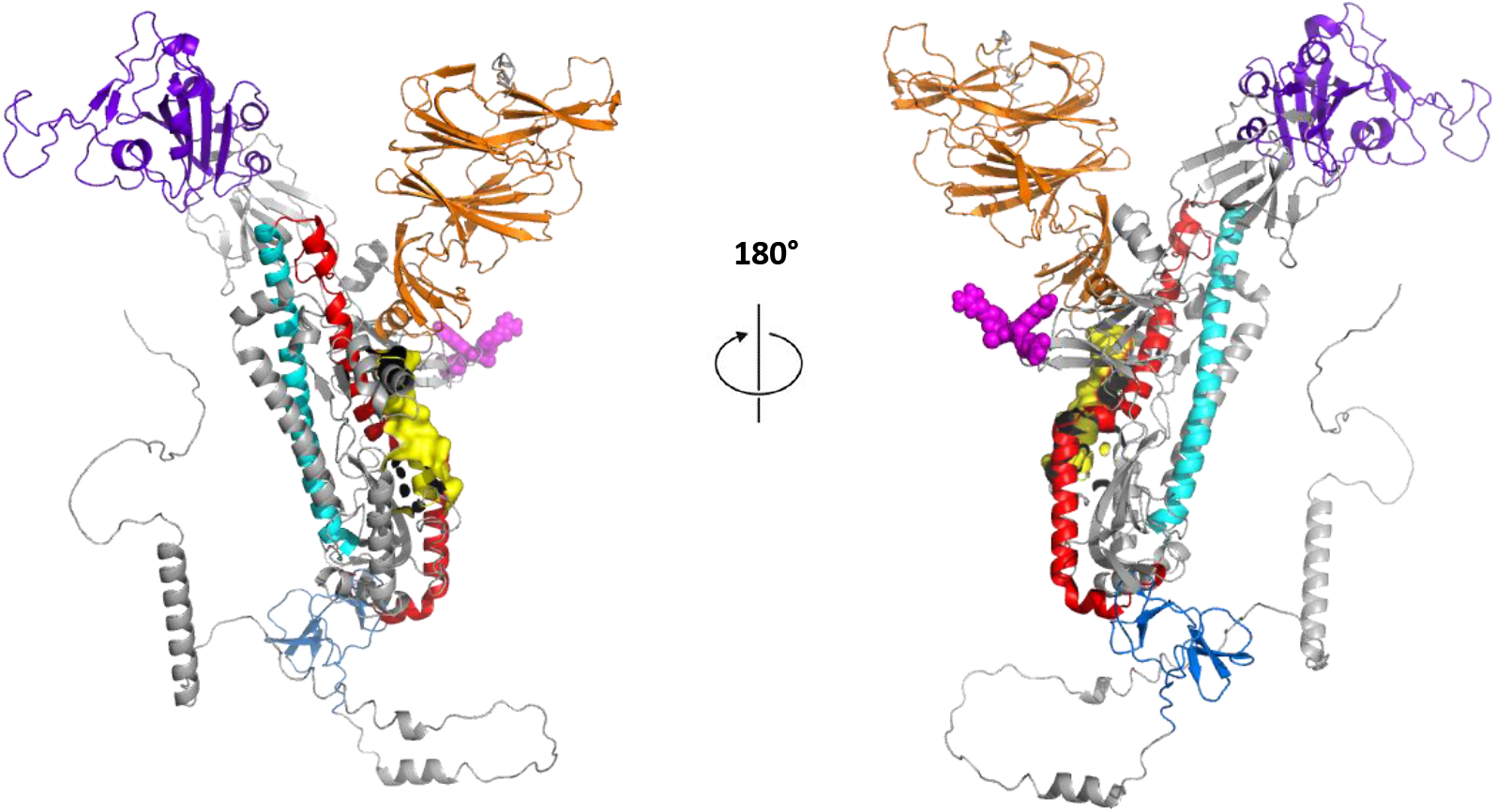
Overall predicted structure for S protein of Wuhan-Hu-1 strain generated by Alphafold. Orange: N-terminal domain, Purple: Receptor binding domain (RBD), Magenta Spheres: S1/S2’ region, Yellow Surface: Fusion protein (FP), Red: heptad repeat, HR1, Cyan: Central helix, and Marine blue: Connector domain.

**Figure 2.**
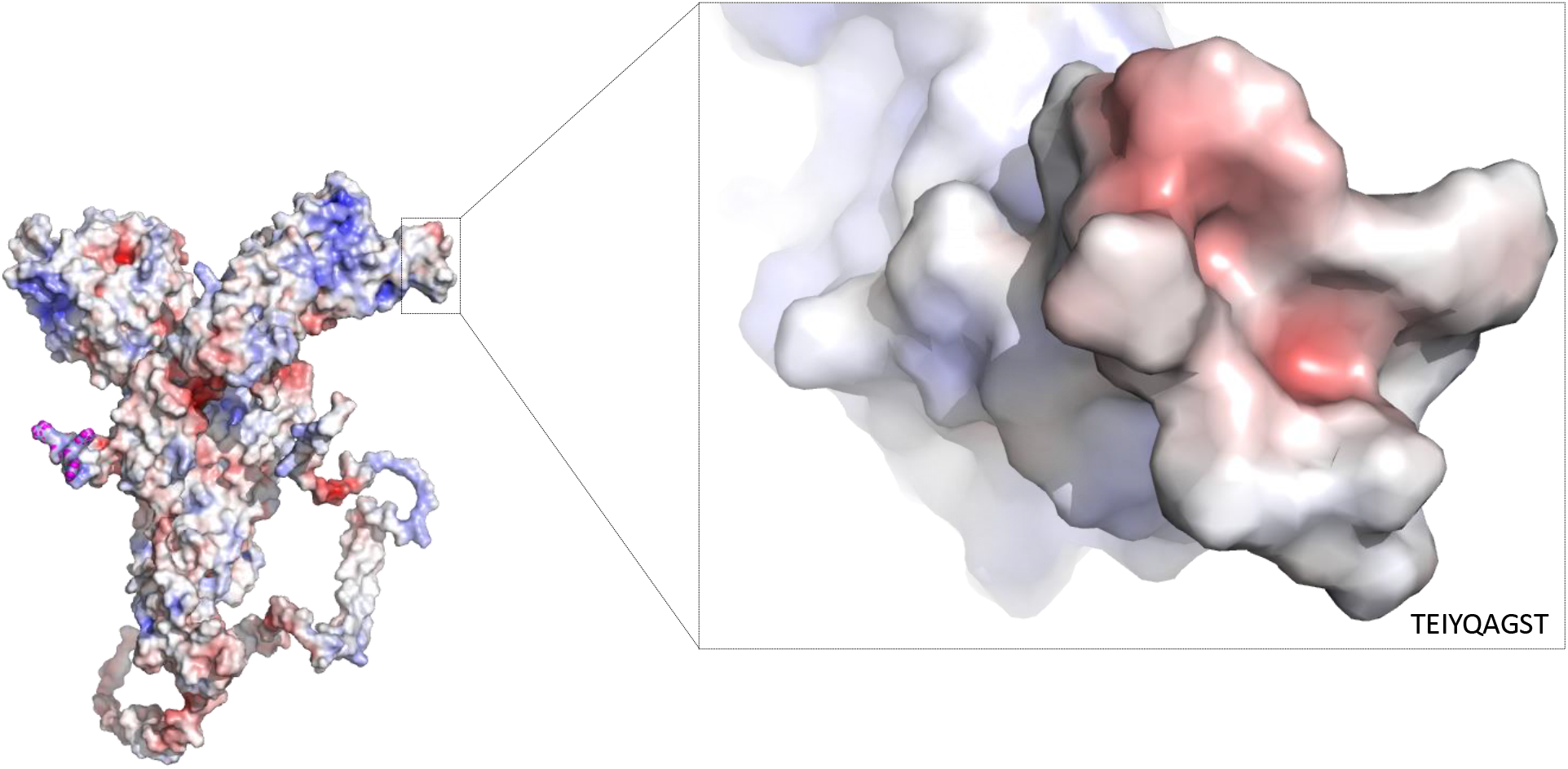
Electrostatic surface for S protein for Wuhan-Hu-1 strain focusing on the recognition site at the RBD for ACE-2 binding showing the high electronegativity generated by Pymol.

**Figure 3.**
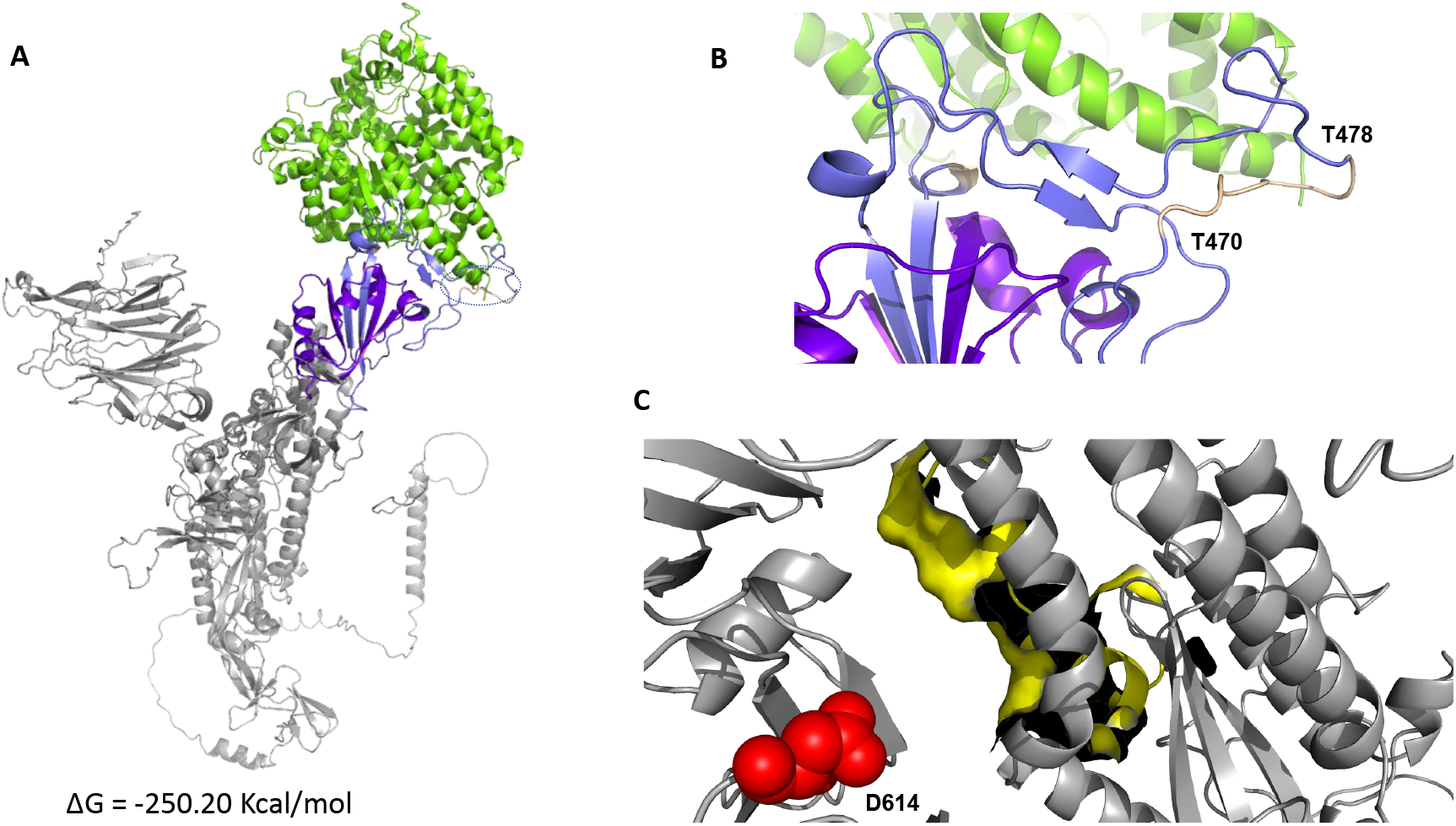
(A) Overall binding mode of ACE-2 (green) towards S protein of Wuhan-Hu-1 strain. (B) Close-up for the binding mode of ACE-2 and recognition site for RBD of S protein. (C) Close-up for D614 along with fusion protein (Yellow surface).

### Structural Characterization of S protein of Delta strain of SARS-CoV-2 and ACE-2

Next, we induced the mutations to elucidate the overall predicted structure of S protein for the Delta variant using alpha fold with an overall of 1271 amino acid (mean pLDDT = 77.71), where the delta variant showed two deleted sequences E156 and F157 (**figure 4**). The electrostatic map showed less electronegativity at the receptor binding domain of S protein where ACE-2 receptor binds, compared to the WT strain (**figure 5**). This was followed by docking of S protein of Delta variant to the ACE-2 receptor extracted from (PDB ID: 6M0J) using Clustpro which showed a change in the RBD of S protein, where two amino acids were mutated L452R and T478K resulting in R450 and K476, respectively at S protein of Delta variant affecting the electrostatic interactions and the recognition sites at the RBD of S protein ΔG = -242.80 Kcal/mol. In addition, the Delta variant exhibited dual mutations close to FB, D164G and D950N resulting in D162 and N948, respectively. This could support the high infection and transmission rates of the Delta variant, where D164G was previously reported for its potential role in promoting the opening of the spike protein to enhance virion spike density and infectivity [30, 31]. In addition, D950N showed proximity to FP that can affect the stability profile of fusion protein leading to a post-fusion state for the whole S protein (**figure 6C**).

**Figure 4.**
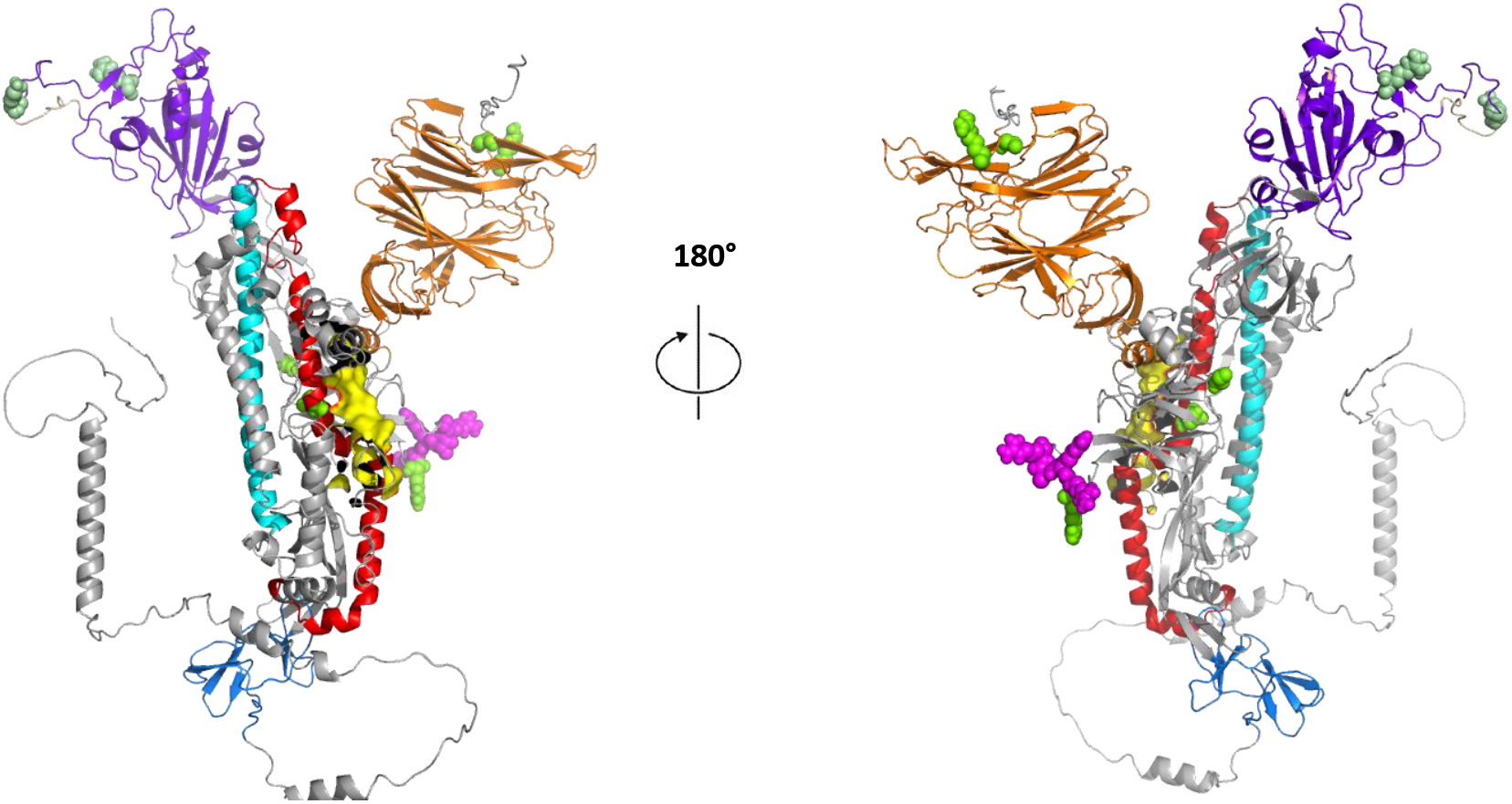
Overall predicted structure for S protein of Delta variant of SARS-CoV-2 generated by Alphafold. Orange: N-terminal domain, Purple: Receptor binding domain (RBD), Magenta Spheres: S1/S2’ region, Yellow Surface: Fusion protein (FP), Red: heptad repeat, HR1, Cyan: Central helix, and Marine blue: Connector domain. Mutations were induced and predicted to be highlighted as green spheres.

**Figure 5.**
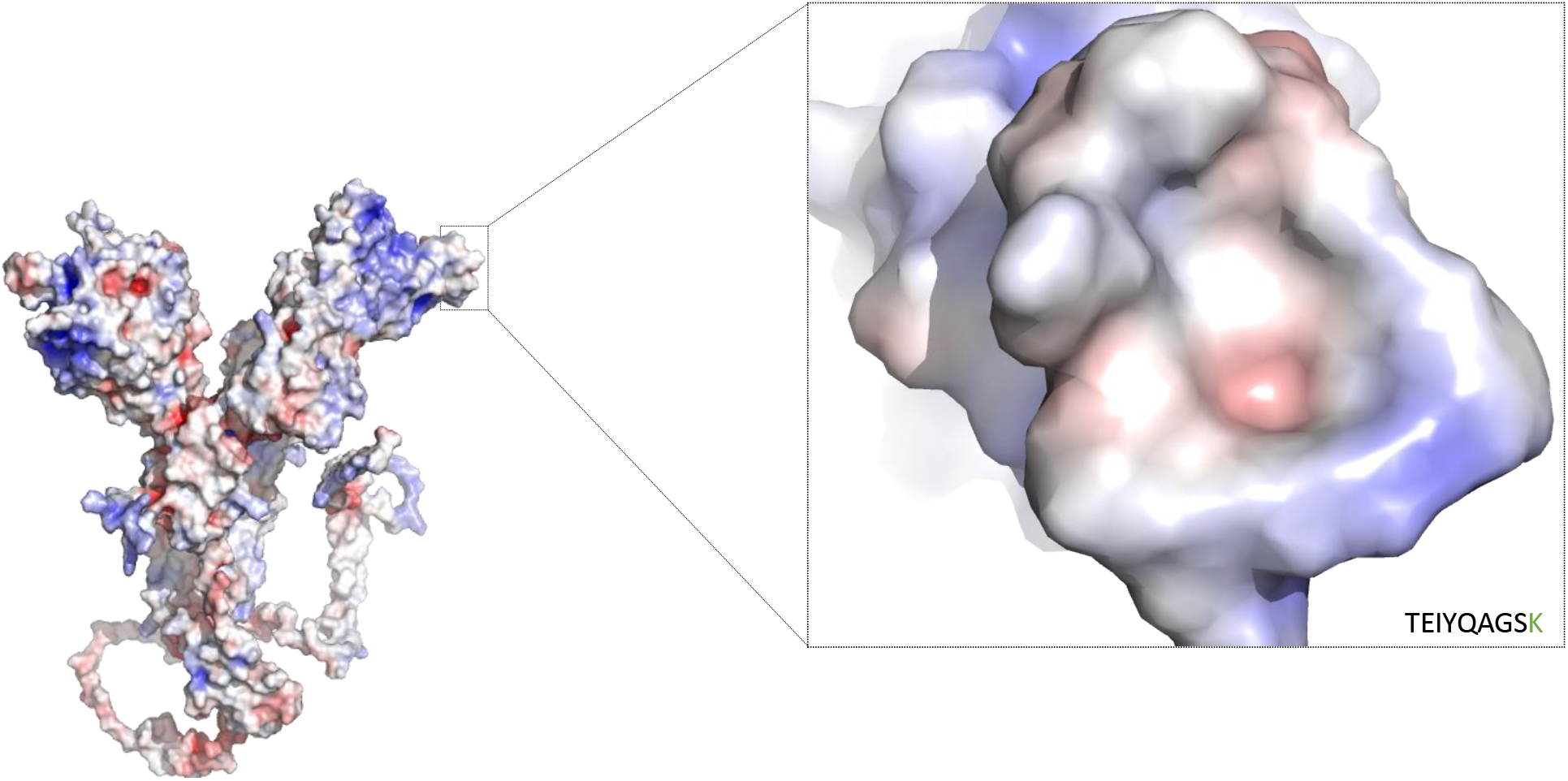
Electrostatic surface for S protein of Delta variant of SARS-CoV-2 focusing on the recognition site at the RBD for ACE-2 binding showing the low electronegativity generated by Pymol.

**Figure 6.**
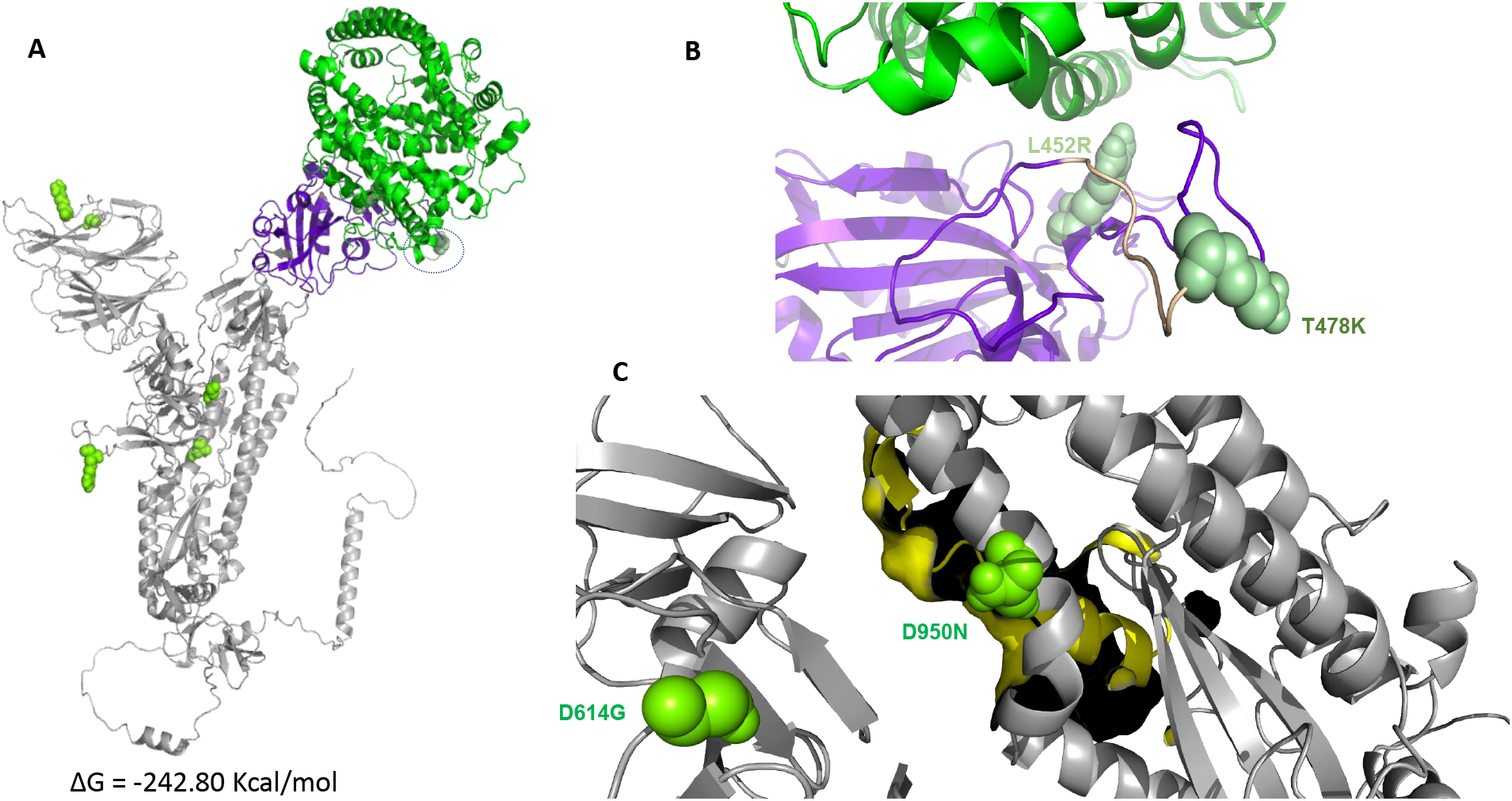
(A) Overall binding mode of ACE-2 (green) towards S protein of Delta variant of SARS-CoV-2. (B) Close-up for the binding mode of ACE-2 and recognition site for RBD of S protein. (C) Close-up for D614G and D950N along with fusion protein (Yellow surface).

### Structural Characterization of S protein of Omicron strain of SARS-CoV-2 and ACE-2

Next, we induced the mutations either by deletion, modification, or insertion to elucidate the overall predicted structure of S protein for the Omicron variant using Alphafold with 1268 amino acids (mean pLDDT= 76.96), where the Omicron variant showed more than 30 mutations at different sites of S protein (**figure 7**). The electrostatic map showed less electronegativity at the receptor binding domain of S protein for Omicron variant where ACE-2 receptor binds, compared to the WT strain (**figure 8**). This was followed by docking of S protein of the Omicron variant to the ACE-2 receptor extracted from PDB (PDB ID: 6M0J) using Clustpro which showed a change in the RBD of S protein, where multiple amino acids were mutated near the recognition site at T478K, S477N, E484A, and Q493A which changed the electrostatic potentials and the recognition sites at the RBD of S protein ΔG = -266.40 Kcal/mol, suggesting a higher binding affinity to ACE-2 receptor. In addition, the Delta variant exhibited a triple mutation in the neighboring amino acid residues near FB, D614G, H655Y, and Q954H. These results support a potential enhanced infectivity and transmission of the Omicron variant mutations by affecting the fusion state of the S protein (**figure 9C**).

**Figure 7.**
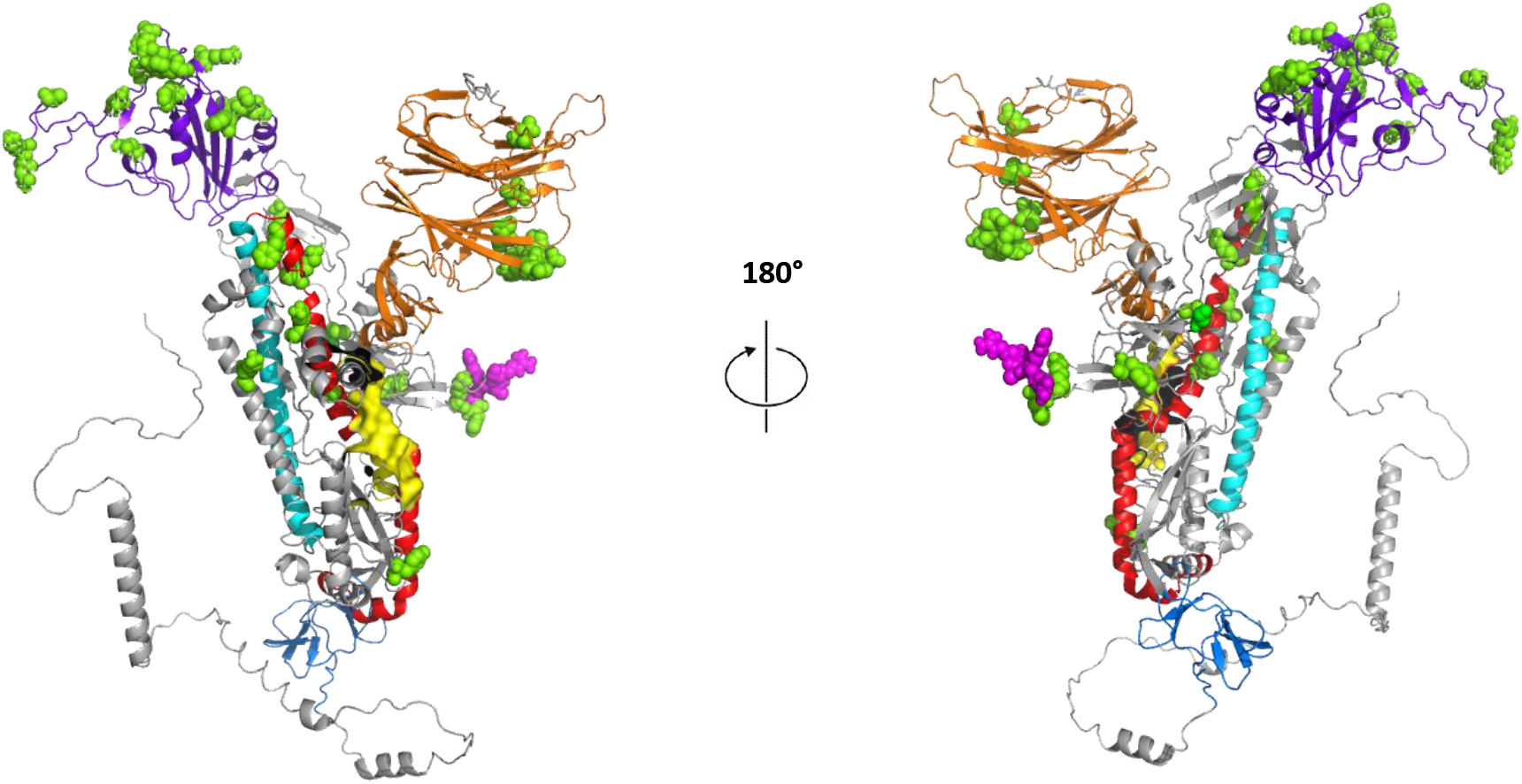
Overall predicted structure for S protein of Omicron variant of SARS-CoV-2 generated by Alphafold. Orange: N-terminal domain, Purple: Receptor binding domain (RBD), Magenta Spheres: S1/S2’ region, Yellow Surface: Fusion protein (FP), Red: heptad repeat, HR1, Cyan: Central helix, and Marine blue: Connector domain. Mutations were induced and predicted to be highlighted as green spheres.

**Figure 8.**
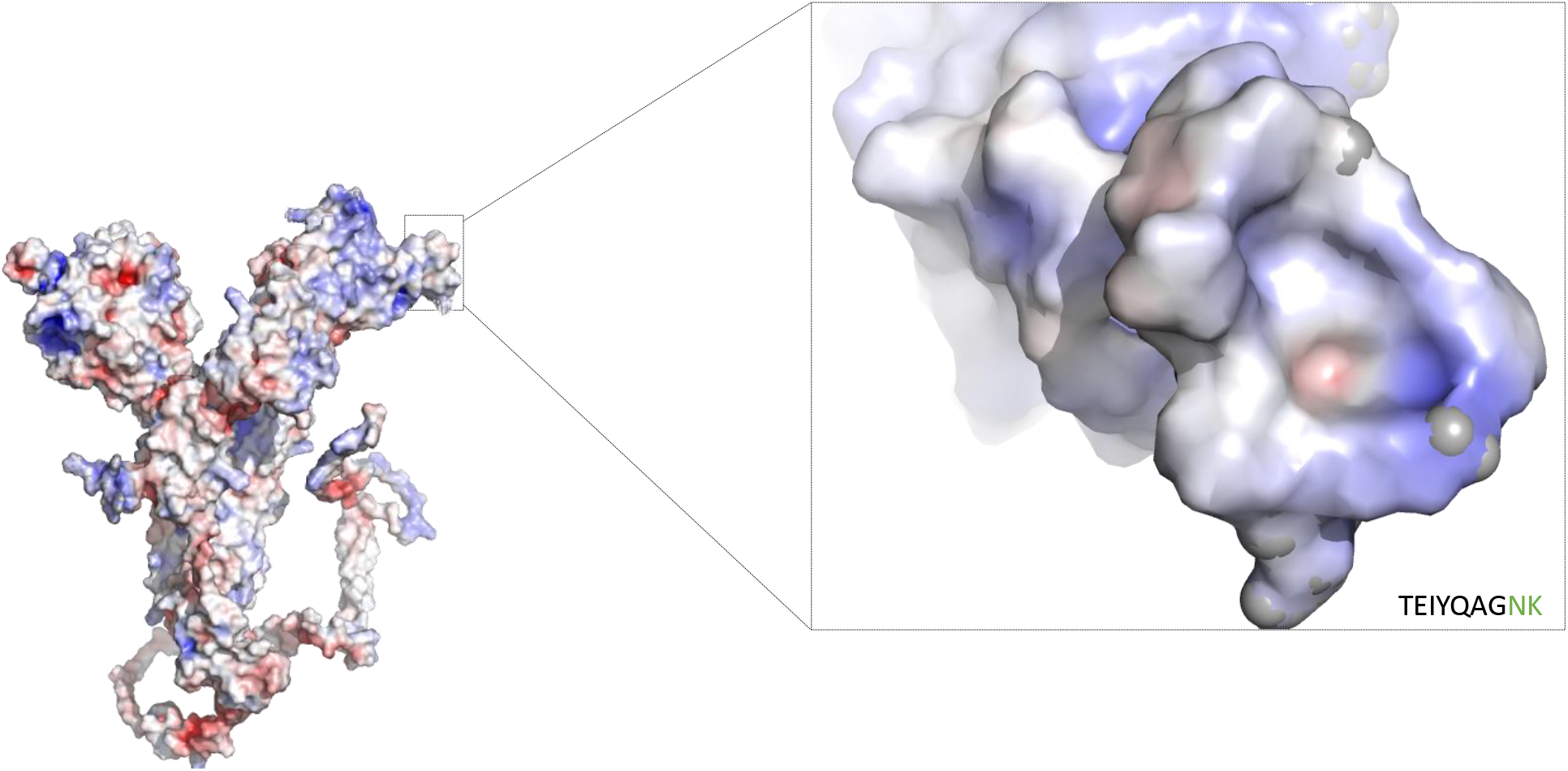
Electrostatic surface for S protein of Omicron variant of SARS-CoV-2 focusing on the recognition site at the RBD for ACE-2 binding showing the low electronegativity generated by Pymol.

**Figure 9.**
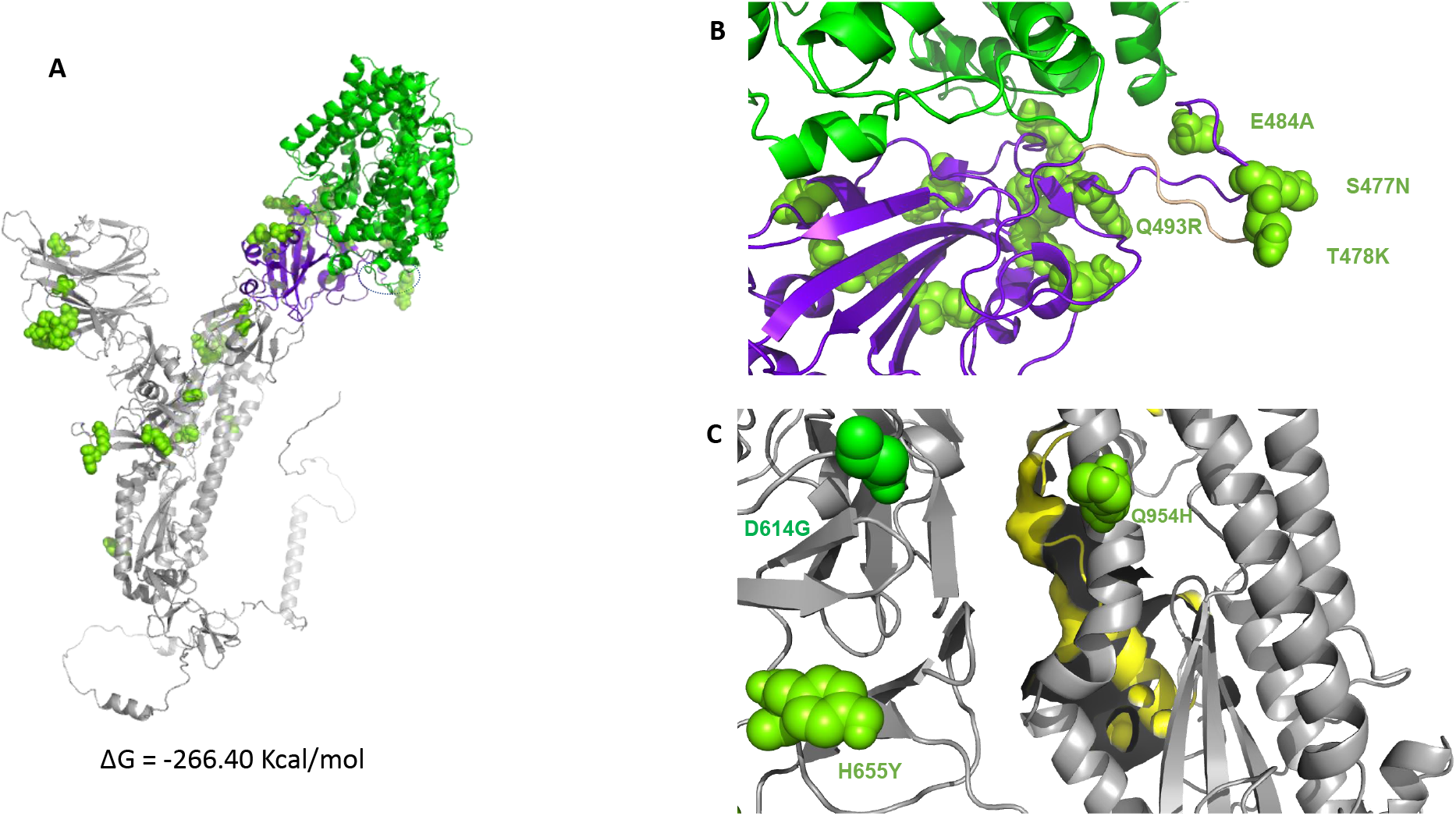
(A) Overall binding mode of ACE-2 (green) towards S protein of Omicron variant of SARS-CoV-2. (B) Close-up for the binding mode of ACE-2 and recognition site for RBD of S protein. (C) Close-up for D614G, H655Y, and Q954H along with fusion protein (Yellow surface).

## Conclusions

In conclusion, we followed an *in silico* computational approach to predict the structures for S protein of Wuhan-Hu-1, Delta, and Omicron variants. We elucidate the significant changes of the RBD, electrostatic potentials for recognition sites at RBD, and FP site using Alphafold. Docking of S protein to the ACE2 receptor demonstrate their energy profiles via electrostatic and Van der Waal scoring functions. This revealed that the ACE2 receptor exhibits an enhanced binding profile towards S protein of Omicron variant, compared to the original WT as well as the Delta variant. In addition, we detected significant changes in the FP sites of S protein for Omicron variant, compared to Wuhan-Hu-1 and Delta variants. Our study suggests that the higher infectivity of the Omicron variant can be explained in part by on the significant mutations in the RBD and the post-fusion enhancement of the FP. Importantly, these results require further validation by X-ray crystallography and/or cryo-EM of the Omicron variant S-protein.

## Supplemental figures and tables

**Figure S1**. Predicted local distance difference test (pLDDT) graphs for predicted S protein for SARS-CoV-2, Delta, and Omicron variants generated by Alphafold.

**Table S1**. Energy profiles for the predicted models for ACE2 bound to S protein of Wuhan-Hu-1 strain.

**Table S2**. Energy profiles for the predicted models for ACE2 bound to S protein of Delta variant.

**Table S3**. Energy profiles for the predicted models for ACE2 bound to S protein of Omicron variant.

