## Supplementary material for "Structural Insights of SARS-CoV-2 Spike Protein from Delta and Omicron Variants": Table S1

| Cluster | Members | Representative | Weighted Score |
|---|---|---|---|
| 0 | 84 | Center | -235.3 |
| 0 | 84 | Lowest Energy | -276 |
| 1 | 75 | Center | -233.9 |
| 1 | 75 | Lowest Energy | -251.7 |
| 2 | 41 | Center | -230 |
| 2 | 41 | Lowest Energy | -233.4 |
| 3 | 38 | Center | -244.1 |
| 3 | 38 | Lowest Energy | -251.4 |
| 4 | 35 | Center | -241.8 |
| 4 | 35 | Lowest Energy | -247 |
| 5 | 30 | Center | -216.6 |
| 5 | 30 | Lowest Energy | -248.9 |
| 6 | 22 | Center | -207.5 |
| 6 | 22 | Lowest Energy | -250.2 |
| 7 | 20 | Center | -210 |
| 7 | 20 | Lowest Energy | -266.4 |
| 8 | 17 | Center | -224.3 |
| 8 | 17 | Lowest Energy | -224.3 |
| 9 | 17 | Center | -207.1 |
| 9 | 17 | Lowest Energy | -216.4 |
| 10 | 16 | Center | -227.4 |
| 10 | 16 | Lowest Energy | -227.4 |
| 11 | 14 | Center | -200.5 |
| 11 | 14 | Lowest Energy | -262.9 |
| 12 | 14 | Center | -210.4 |
| 12 | 14 | Lowest Energy | -241.3 |
| 13 | 13 | Center | -207.7 |
| 13 | 13 | Lowest Energy | -238.7 |
| 14 | 13 | Center | -219.3 |
| 14 | 13 | Lowest Energy | -219.3 |
| 15 | 13 | Center | -206 |
| 15 | 13 | Lowest Energy | -221.1 |
| 16 | 12 | Center | -259.9 |
| 16 | 12 | Lowest Energy | -259.9 |
| 17 | 12 | Center | -204.8 |
| 17 | 12 | Lowest Energy | -217.8 |
| 18 | 11 | Center | -204.1 |
| 18 | 11 | Lowest Energy | -221.9 |
| 19 | 10 | Center | -205.8 |
| 19 | 10 | Lowest Energy | -211.8 |
| 20 | 10 | Center | -217.8 |
| 20 | 10 | Lowest Energy | -217.8 |
| 21 | 10 | Center | -211.7 |
| 21 | 10 | Lowest Energy | -211.7 |
| 22 | 10 | Center | -200.1 |
| 22 | 10 | Lowest Energy | -208.7 |
| 23 | 7 | Center | -221.3 |
| 23 | 7 | Lowest Energy | -224.8 |
| 24 | 7 | Center | -207.5 |
| 24 | 7 | Lowest Energy | -240.3 |
| 25 | 6 | Center | -204.7 |
| 25 | 6 | Lowest Energy | -231.2 |
| 26 | 6 | Center | -238.3 |
| 26 | 6 | Lowest Energy | -238.3 |
| 27 | 5 | Center | -201.1 |
| 27 | 5 | Lowest Energy | -211.2 |
| 28 | 3 | Center | -196.9 |
| 28 | 3 | Lowest Energy | -203.2 |
