## Supplementary material for "Structural Insights of SARS-CoV-2 Spike Protein from Delta and Omicron Variants": Table S2

| Cluster | Members | Representative | Weighted Score |
|---|---|---|---|
| 0 | 65 | Center | -253.5 |
| 0 | 65 | Lowest Energy | -257.4 |
| 1 | 55 | Center | -248.6 |
| 1 | 55 | Lowest Energy | -265.5 |
| 2 | 46 | Center | -209.2 |
| 2 | 46 | Lowest Energy | -242.8 |
| 3 | 36 | Center | -220.3 |
| 3 | 36 | Lowest Energy | -229 |
| 4 | 30 | Center | -237.7 |
| 4 | 30 | Lowest Energy | -264.8 |
| 5 | 29 | Center | -220.8 |
| 5 | 29 | Lowest Energy | -228.7 |
| 6 | 28 | Center | -233.8 |
| 6 | 28 | Lowest Energy | -260.1 |
| 7 | 26 | Center | -208.6 |
| 7 | 26 | Lowest Energy | -261.9 |
| 8 | 24 | Center | -211.4 |
| 8 | 24 | Lowest Energy | -229.7 |
| 9 | 23 | Center | -226.8 |
| 9 | 23 | Lowest Energy | -261.5 |
| 10 | 17 | Center | -233.6 |
| 10 | 17 | Lowest Energy | -233.6 |
| 11 | 14 | Center | -215.8 |
| 11 | 14 | Lowest Energy | -231 |
| 12 | 14 | Center | -213.3 |
| 12 | 14 | Lowest Energy | -223.2 |
| 13 | 13 | Center | -212 |
| 13 | 13 | Lowest Energy | -231.9 |
| 14 | 13 | Center | -217.3 |
| 14 | 13 | Lowest Energy | -228.4 |
| 15 | 13 | Center | -215.2 |
| 15 | 13 | Lowest Energy | -228.1 |
| 16 | 12 | Center | -223.6 |
| 16 | 12 | Lowest Energy | -230.9 |
| 17 | 12 | Center | -204.8 |
| 17 | 12 | Lowest Energy | -251.5 |
| 18 | 12 | Center | -249.4 |
| 18 | 12 | Lowest Energy | -249.4 |
| 19 | 12 | Center | -237.8 |
| 19 | 12 | Lowest Energy | -237.8 |
| 20 | 12 | Center | -229.2 |
| 20 | 12 | Lowest Energy | -229.2 |
| 21 | 11 | Center | -226.5 |
| 21 | 11 | Lowest Energy | -226.5 |
| 22 | 11 | Center | -214.6 |
| 22 | 11 | Lowest Energy | -241.4 |
| 23 | 10 | Center | -215 |
| 23 | 10 | Lowest Energy | -243.8 |
| 24 | 10 | Center | -250.4 |
| 24 | 10 | Lowest Energy | -250.4 |
| 25 | 10 | Center | -207.9 |
| 25 | 10 | Lowest Energy | -219.1 |
| 26 | 9 | Center | -208.8 |
| 26 | 9 | Lowest Energy | -240.2 |
| 27 | 6 | Center | -208.8 |
| 27 | 6 | Lowest Energy | -252.4 |
| 28 | 6 | Center | -208.3 |
| 28 | 6 | Lowest Energy | -218 |
| 29 | 6 | Center | -234.8 |
| 29 | 6 | Lowest Energy | -234.8 |
