## Supplementary material for "Structural Insights of SARS-CoV-2 Spike Protein from Delta and Omicron Variants": Table S3

| Cluster | Members | Representative | Weighted Score |
|---|---|---|---|
| 0 | 85 | Center | -208 |
| 0 | 85 | Lowest Energy | -285.4 |
| 1 | 53 | Center | -212.2 |
| 1 | 53 | Lowest Energy | -237.7 |
| 2 | 29 | Center | -201.9 |
| 2 | 29 | Lowest Energy | -266.4 |
| 3 | 29 | Center | -237.3 |
| 3 | 29 | Lowest Energy | -237.3 |
| 4 | 28 | Center | -239.2 |
| 4 | 28 | Lowest Energy | -239.3 |
| 5 | 27 | Center | -210.2 |
| 5 | 27 | Lowest Energy | -230.6 |
| 6 | 25 | Center | -214.8 |
| 6 | 25 | Lowest Energy | -214.8 |
| 7 | 25 | Center | -211.4 |
| 7 | 25 | Lowest Energy | -217.5 |
| 8 | 20 | Center | -225 |
| 8 | 20 | Lowest Energy | -225 |
| 9 | 20 | Center | -212.7 |
| 9 | 20 | Lowest Energy | -228.5 |
| 10 | 19 | Center | -231.1 |
| 10 | 19 | Lowest Energy | -231.1 |
| 11 | 17 | Center | -235.8 |
| 11 | 17 | Lowest Energy | -235.8 |
| 12 | 16 | Center | -201.1 |
| 12 | 16 | Lowest Energy | -228.4 |
| 13 | 15 | Center | -234.9 |
| 13 | 15 | Lowest Energy | -234.9 |
| 14 | 14 | Center | -200.8 |
| 14 | 14 | Lowest Energy | -221.5 |
| 15 | 13 | Center | -205 |
| 15 | 13 | Lowest Energy | -216.5 |
| 16 | 12 | Center | -264 |
| 16 | 12 | Lowest Energy | -264 |
| 17 | 12 | Center | -220.9 |
| 17 | 12 | Lowest Energy | -223.1 |
| 18 | 11 | Center | -202.1 |
| 18 | 11 | Lowest Energy | -267.7 |
| 19 | 11 | Center | -204.7 |
| 19 | 11 | Lowest Energy | -235 |
| 20 | 11 | Center | -206.2 |
| 20 | 11 | Lowest Energy | -226.9 |
| 21 | 11 | Center | -210 |
| 21 | 11 | Lowest Energy | -231.5 |
| 22 | 10 | Center | -205.8 |
| 22 | 10 | Lowest Energy | -247.6 |
| 23 | 10 | Center | -256.4 |
| 23 | 10 | Lowest Energy | -256.4 |
| 24 | 10 | Center | -227.2 |
| 24 | 10 | Lowest Energy | -227.2 |
| 25 | 10 | Center | -226.3 |
| 25 | 10 | Lowest Energy | -242 |
| 26 | 10 | Center | -223.4 |
| 26 | 10 | Lowest Energy | -233.3 |
| 27 | 9 | Center | -202.1 |
| 27 | 9 | Lowest Energy | -213.8 |
| 28 | 7 | Center | -205.8 |
| 28 | 7 | Lowest Energy | -222.9 |
| 29 | 6 | Center | -210.2 |
| 29 | 6 | Lowest Energy | -212.3 |
