## Supplementary figures and images for "Structural Insights of SARS-CoV-2 Spike Protein from Delta and Omicron Variants"

### Figure S1

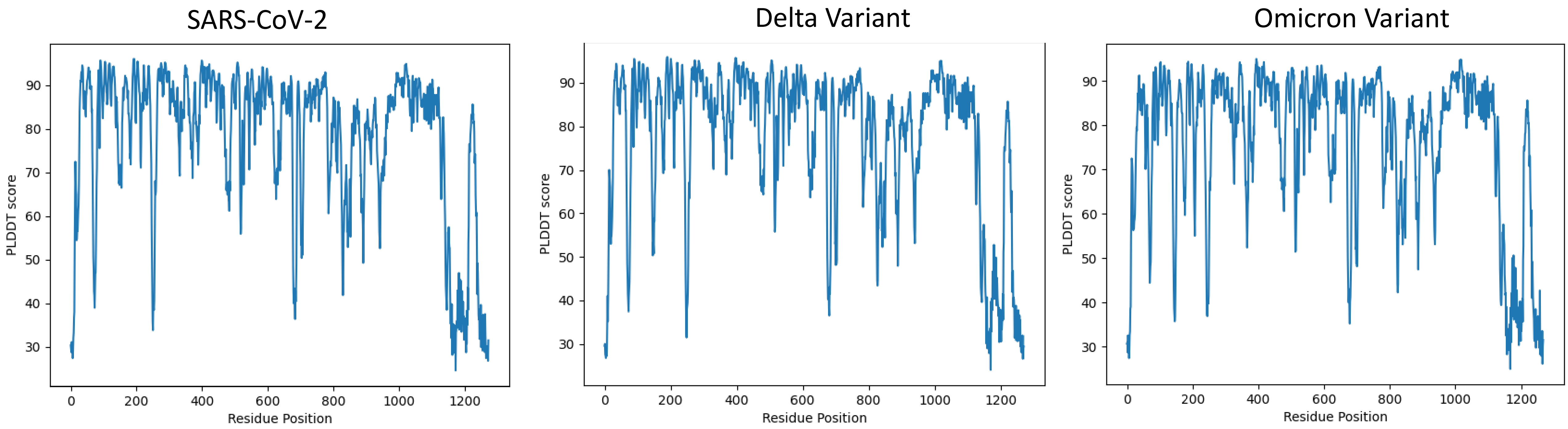
